# Benchmarking small-variant genotyping in polyploids

**DOI:** 10.1101/2021.03.29.436766

**Authors:** Daniel P Cooke, David C Wedge, Gerton Lunter

**Affiliations:** MRC Weatherall Institute of Molecular Medicine, University of Oxford, Oxford, UK; Manchester Cancer Research Centre, University of Manchester, Manchester, UK; Department of Epidemiology, University Medical Centre Groningen, Groningen, The Netherlands

**Author notes:** Correspondence should be addressed to D.P.C.

## Abstract

Genotyping from sequencing is the basis of emerging strategies in the molecular breeding of polyploid plants. However, compared with the situation for diploids, where genotyping accuracies are confidently determined with comprehensive benchmarks, polyploids have been neglected; there are no benchmarks measuring genotyping error rates for small variants using real sequencing reads. We previously introduced a variant calling method – Octopus – that accurately calls germline variants in diploids and somatic mutations in tumors. Here, we evaluate Octopus and other popular tools on whole-genome tetraploid and hexaploid datasets created using *in silico* mixtures of diploid Genome In a Bottle samples. We find that genotyping errors are abundant for typical sequencing depths, but that Octopus makes 25% fewer errors than other methods on average. We supplement our benchmarks with concordance analysis in real autotriploid banana datasets.

---

Polyploidy is common in many plant species, including important agricultural crops such as wheat, potato, oat, coffee, rapeseed, cotton, banana, and sugar cane^1^. In mammals, polyploidization regularly occurs during tumorigenesis, but has also been shown to be a normal part of development in some mouse and humans tissues^2^. Molecular markers have been widely used for decades in artificial polyploid crop breeding to assist selection of more desirable traits such as better resilience to climate change and disease. More recently, genotyping by sequencing has been applied for marker assisted and genomic selection^3–5^, and the assembly of high-quality plant reference genomes^6–10^, together with developments in resequencing, promise new strategies for quantitative trait analysis with a wider variety of genetic variants and better linkage information than is currently possible^10–13^.

Despite these advances, methods for genotyping polyploids from sequencing data has received little scrutiny in comparison to those for diploids^13–16^. Variant calling and genotyping in polyploids is more difficult than in diploids primarily because the number of possible genotypes at a given loci is combinatorial in the ploidy and number of distinct alleles, and a sequencing read cannot distinguish identical allele copies in the absence of physical linkage with other heterozygous alleles. It therefore becomes harder to determine the copy number of a particular allele for a fixed read depth as the ploidy increases. The lower per-allele coverage also makes it harder to distinguish sequencing error from true variants. Haplotype-based methods increase power to genotype individual alleles by jointly evaluating combinations of several proximal alleles (haplotypes). They are now standard for diploid calling^17–22^, and are becoming more common for somatic mutation calling in tumours^17^. Unfortunately, only a minority are capable of polyploid calling^17–19^, and none have been rigorously tested for this purpose. Specialised methods for polyploid genotyping have been developed^23–25^, but are only suitable for biallelic SNPs. Crucially, existing benchmarks of polyploid calling methods fall short of the standard demanded for diploid calling^12,16,26,27^. In particular, we are not aware of any that consider indels, genotyping errors in real sequencing data, or representation differences between callers^16^. Polyploid genotyping error rates from sequencing are therefore highly uncertain, undermining developments that depends on them.

We sought to address some of these issues by conducting an in-depth assessment of polyploid small variant calling using an independent and comprehensive ground truth, real sequencing data, and haplotype-aware comparisons. Our analysis, implemented in Snakemake^28^, is made available online at https://github.com/luntergroup/polyploid.

## Results

### Synthetic polyploid genomes

We created synthetic tetraploid and hexaploid samples with high quality truth sets by merging Genome In A Bottle^14^ (GIAB) v4.2 GRCh38 variants for human diploid samples HG002, HG003, and HG004. We chose HG003 and HG004 for the tetraploid sample – the two unrelated parents of HG002. Evaluation regions were defined by intersecting^29^ the GIAB high confidence regions for each sample, resulting in 2.50Gb (86% non-N primary reference) confident tetraploid bases containing 5, 095.314 variants, and 2.49Gb (85% non-N primary reference) confident hexaploid bases containing 5, 028, 566 variants. We constructed polyploid Illumina NovaSeq whole-genome test data by mixing reads generated independently for each sample with consistent PCR-free library preparation and depths (Methods). Each individual sequencing run targeted 35x coverage, resulting in 70x coverage tetraploid samples and 105x coverage hexaploid samples. We confirmed total read counts were similar for each contributing sample, ensuring realistic heterozygous allele frequencies. We then randomly downsampled the full datasets, starting from 10x in 10x intervals to the full coverage, resulting in 6 + 10 = 16 polyploid datasets. All reads were mapped to GRCh38 with BWA-MEM^30^ (Methods).

### Polyploid genotyping accuracy from short-read WGS

We evaluated three popular germline variant callers that support polyploid genotypes: Octopus^17^; GATK4^18^; and FreeBayes^19^, on all synthetic polyploid Illumina datasets, and in the diploid HG002 sample to get performance baselines. Other notable germline callers, such as DeepVariant^21^, Strelka2^20^, and Platypus^22^, were not included because they do not support polyploid calling. We also ignored methods that call polyploid SNVs but not indels, such as polyRAD^24^. Other than specifying the ploidy and requesting genotype qualities from FreeBayes, we used default setting for all callers (Methods). Octopus calls were hard-filtered with default thresholds, GATK4 and FreeBayes calls were hard-filtered using recommended thresholds (Methods). Variants were compared using RTG Tools vcfeval_31_ on the basis of both genotype and allele matches (Methods).

Genotyping accuracy was considerably worse for polyploids compared with diploids (Fig. 1a and Supplementary Table 1). For 30x sequencing depth, on average 1/200 diploid genotype calls were incorrect, in contrast with 1/11 for tetraploid and 1/6 for hexaploid. Sensitivity was similarly affected; there were 8x and 16x more false negatives on average for tetraploid and hexaploid, respectively, compared with diploid, for 30x sequencing. There were also more substantial differences in accuracy between callers for polyploids compared with diploids. Notably, Octopus made 26% less errors than GATK4 and 30% less errors than FreeBayes overall, and half the errors on some depths; the largest F-measure difference between callers occurred at moderate sequencing depth: 30x for tetraploid, 50.*x* for hexaploid. F-measure showed a typical logarithmic relationship with sequencing depth for both tetraploid and hexaploid samples, but also suboptimal response considering ploidy; the F-measure lost from doubling the ploidy was not recovered by doubling the depth, and the differential increased with depth.

**Fig. 1.**
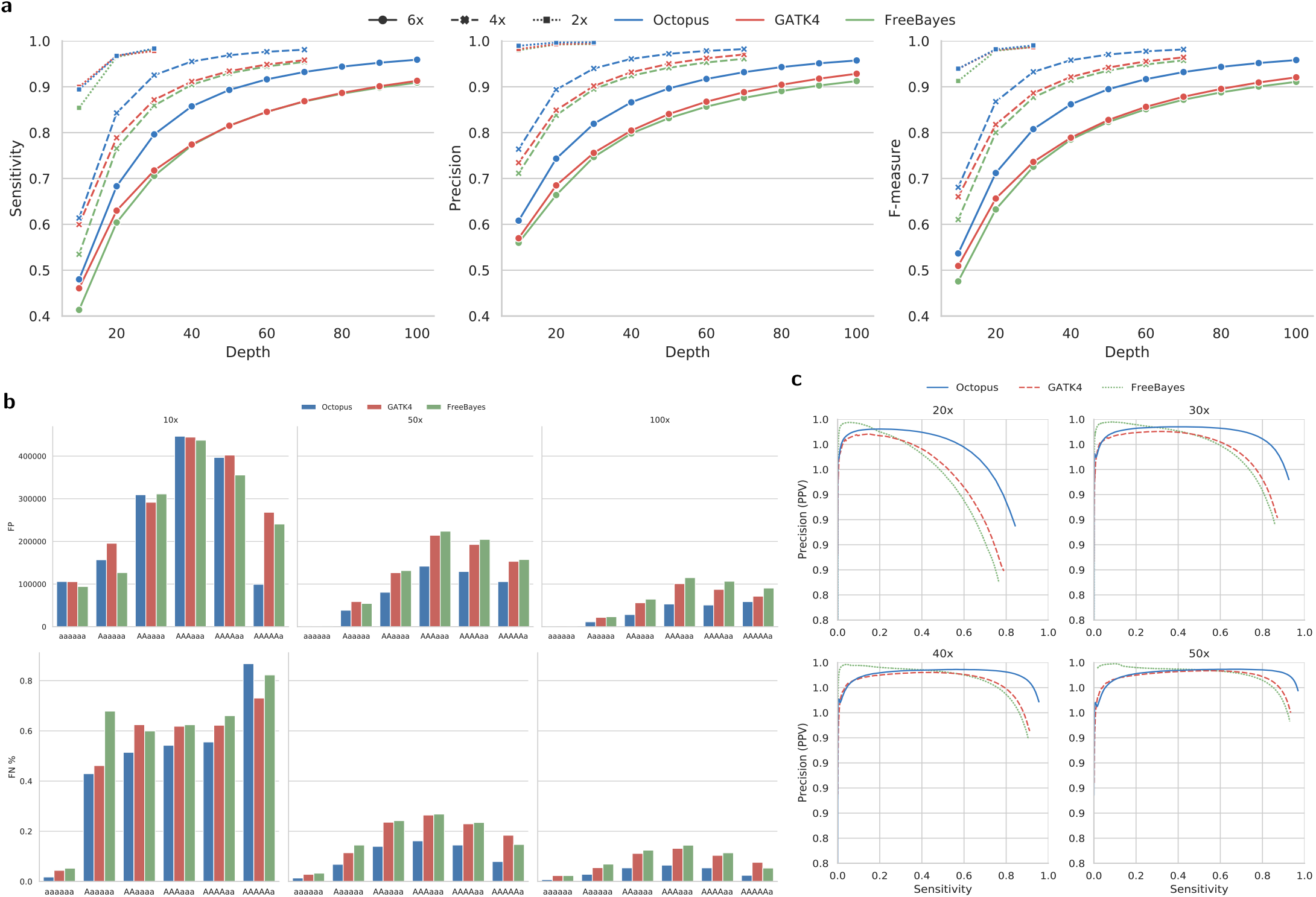
Genotyping accuracy in synthetic polyploids. **a** Sensitivity and precision by depth for each caller on real diploid (2x), and synthetic tetraploid (4x) and hexaploid (6x) Illumina datasets. **b** Counts of false positive biallelic genotypes stratified by depth and copy-number (top). Proportion of false negative biallelic genotypes stratified by depth and copy-number (bottom). **c** Precision-recall curves for a various tetraploid sequencing depths. Score metrics used to generate the curves were RFGQ (Octopus), GQ (GATK4), and GQ (FreeBayes).

The majority of false positives resulted from incorrect genotype copy-numbers: 89% of false positive biallelic genotype calls (97% of all false positives) were due to incorrect copy number. The most common false positive for all depths were the balanced heterozygotes: AAaa and AAAaaa (Fig. 1b, Supplementary Fig. 1, and Supplementary Table 2), 94% of which were due to incorrect copy number. A larger fraction of these were made when the true genotype had —1 alternative allele copy rather than +1 copy (65% vs 34%; Supplementary Fig’s. 2 and 3, and Supplementary Table 3). The most common biallelic false negatives in tetraploids were simplex heterozygotes (those with a single variant copy), while for hexaploids it was duplex heterozygotes (Supplementary Fig’s. 2 and 3). However, normalising by the true prevalence shows that the most frequent false negative for depths ≥ 30x is the balanced heterozygote for both tetraploid and hexaploid; for depths ≤ 20x the most frequent false negative was simplex (Fig. 1b). Interestingly, there was a slight tendency to miss-call balanced heterzygotes —1 alternative allele copy rather than +1 copy for all callers (Supplementary Fig’s. 2 and 3).

Genotype quality scores were generally well calibrated for all callers (Fig. 1c and Supplementary Fig. 4). However, filtering did not always improve F-measure; the average F-measure percentage change for filtered verses unfiltered calls on all tests was —0.1%, —0.2%, and +3.5%, for Octopus, GATK4, and FreeBayes, respectively. Performance differentials between callers were similar for unfiltered calls (Supplementary Table 1), showing that most of Octopus’ performance advantage come from better genotyping rather than filtering.

Comparison on the basis of allele matching showed less performance differential between callers, ploidies, and depths, particularly for precision (Supplementary Fig./ 5 and Supplementary Table 1). However, predominantly due to better sensitivity at low depths, Octopus still made considerably fewer errors in total than GATK (16% fewer) and FreeBayes (36% fewer).

### Longer haplotypes improve genotyping accuracy

A plausible explanation for Octopus having better genotyping accuracy than GATK4 and FreeBayes is that Octopus considered longer haplotypes – on average – when calculating genotype likelihoods. If the true set of haplotypes including a subset of variants can be confidently determined, then the variance in the genotype posterior probability distribution is expected to decrease, in particularly with respect to copy number, with larger subsets (and therefore longer haplotypes) since the number of discriminating reads is expected to be proportional to the haplotype length (Fig. 2). To test this, we recalled genotypes in the 30x tetraploid sample using a parametrisation of Octopus designed to generate longer haplotypes than with default settings (Methods). The mean called haplotype length increased from 320 bases to 511 bases and the number of false positives decreased by 31,921, but the number of false negatives increased by 26,533.

**Fig. 2.**
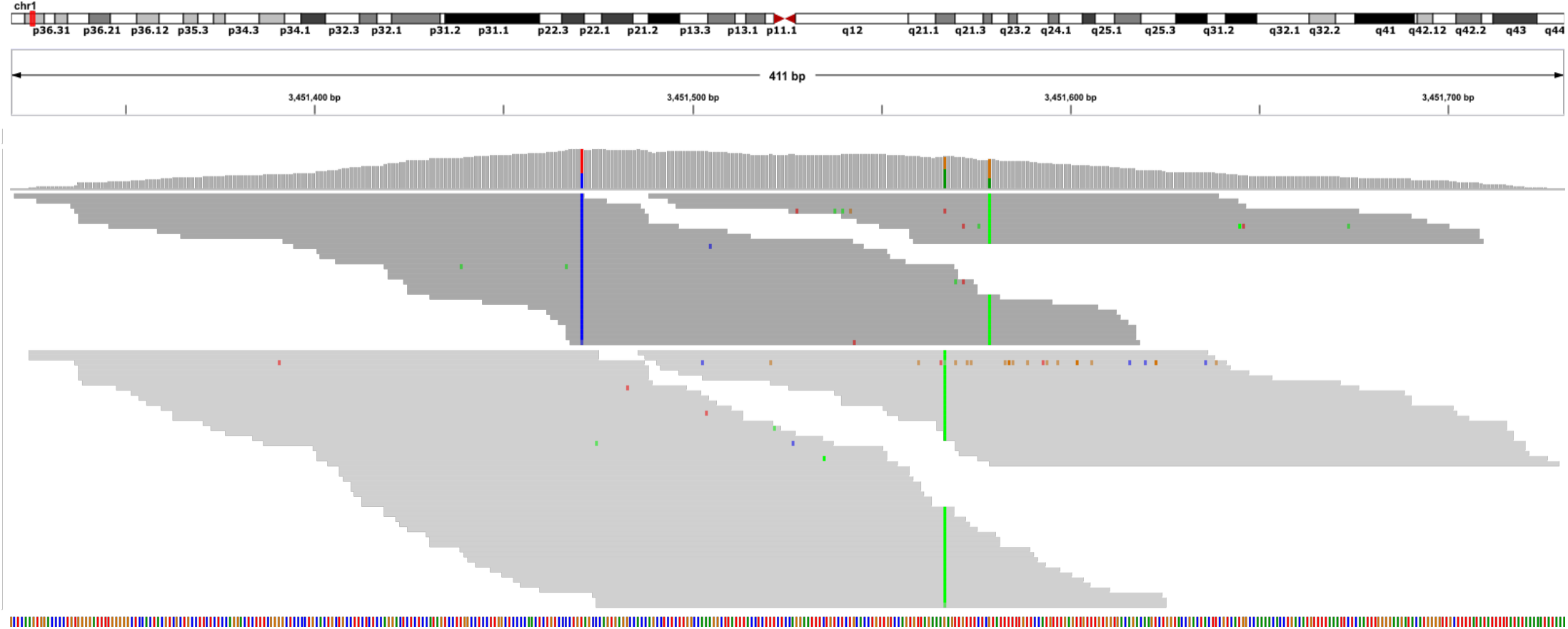
Read pileup of HG003-HG004 tetraploid colored and grouped by supported haplotype. There are two distinct haplotypes (light and dark grey). The true genotypes for the three SNVs (T>C, G>A, G>A) are AAAa, Aaaa, AAAa. The alternative allele read depths are 30/78 (38%), 38/64 (59%), and 20/58 (34%) respectively. GATK4 and FreeBayes both miscall the first two SNVs as AAaa – the most likely genotypes assuming binomially distributed allele observations. Octopus makes the correct calls because it phases all three SNVs, and the first haplotype (including the first and third SNVs) is supported by 74/114 (65%) of reads.

### Banana genotyping

Dwarf Cavendish banana (*Musa acuminata*) is autotriploid consisting of 11 chromosomes with a haploid genome size of around 523Mb, and is an important food source and export-product for many developing countries^7^. To support our previous results on real polyploid samples, we called variants (Methods) in a Dwarf Cavendish banana specimen that was previously whole-genome sequenced with two Illumina technologies, NextSeq-500 and HiSeq-1500, to 65x and 55x coverage, respectively^32^. Both datasets were mapped to the DH Pahang v2 reference^7^ with BWA-MEM, and genotypes were called with Octopus, GATK4, and FreeBayes.

Due to lack of truth data, we evaluated concordance on the two banana datasets using haplotype-aware intersections (Methods). Genotypes called by all callers in both datasets, while substantially the largest intersection set, only accounted for 41% of all distinct genotype calls; 20% of calls were unique to a single callset (Fig. 3). However, there were considerable differences in concordance between the two datasets for each caller: GATK4 had 40% more discordant calls compared with Octopus and 18% more than FreeBayes, despite making 0.6% less calls overall than FreeBayes and only 3% more than Oc-topus (Table 1). We also found high disconcordance when intersecting by called alleles (Table 1 and Supplementary Fig. 6); only 57% of distinct alleles were present in all callsets while 11% were unique to a single callset, indicating that that, in comparison to our results on synthetic data, a larger proportion of false calls arise from incorrect variant alleles rather than copy-number errors.

**Fig. 3.**
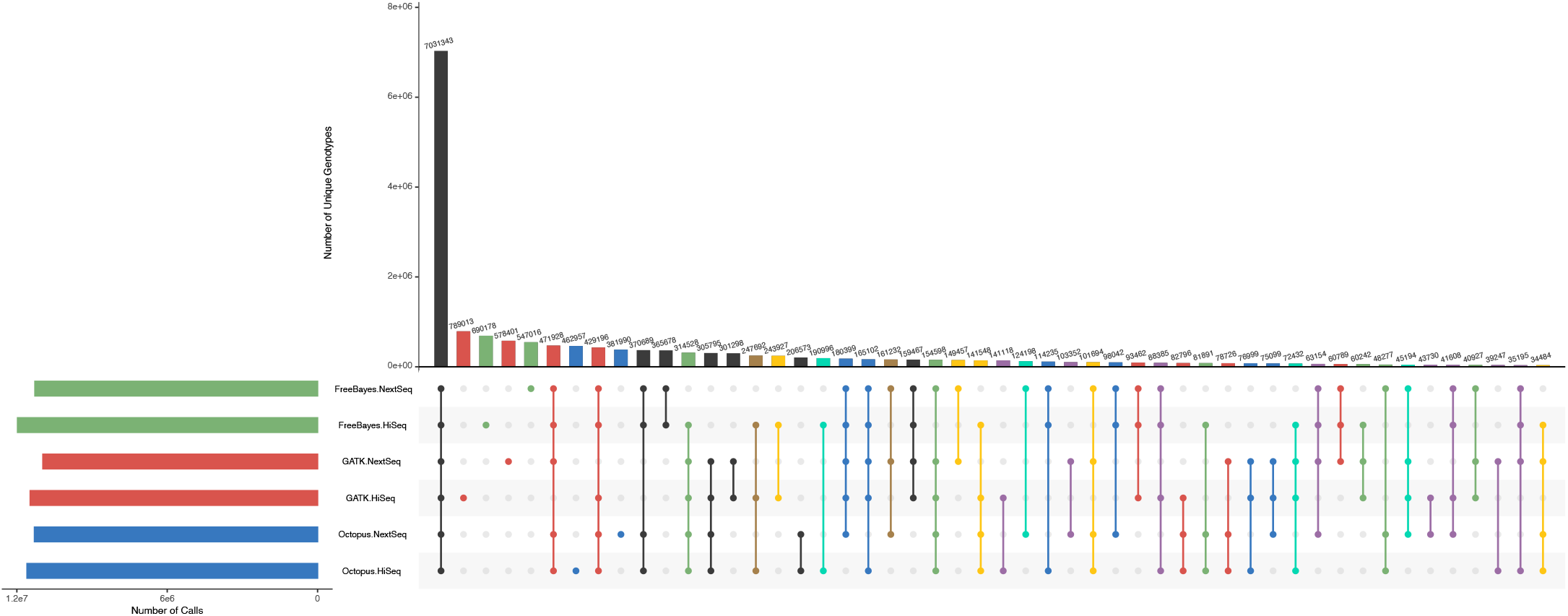
Comparison of genotypes called in two Illumina datasets (HiSeq and NextSeq) of banana specimen by Octopus, GATK4, and FreeBayes. UpSet plot shows callset intersections for each caller-dataset pair. The largest 50/63 intersection sets are shown. Intersections are color coded by caller discordance between the two datasets: No discordances (black), Octopus (blue), GATK4 (red), FreeBayes (green), Octopus & GATK4 (purple), Octopus & FreeBayes (cyan), GATK4 & FreeBayes (yellow), All (brown). The total number of unique genotype calls was 17,151,421.

**Table 1.**
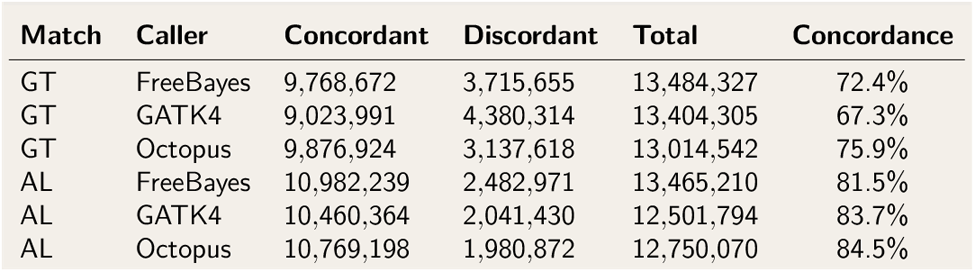
Concordance in two banana Illumina datasets

| Match | Caller | Concordant | Discordant | Total | Concordance |
| --- | --- | --- | --- | --- | --- |
| GT | FreeBayes | 9,768,672 | 3,715,655 | 13,484,327 | 72.4% |
| GT | GATK4 | 9,023,991 | 4,380,314 | 13,404,305 | 67.3% |
| GT | Octopus | 9,876,924 | 3,137,618 | 13,014,542 | 75.9% |
| AL | FreeBayes | 10,982,239 | 2,482,971 | 13,465,210 | 81.5% |
| AL | GATK4 | 10,460,364 | 2,041,430 | 12,501,794 | 83.7% |
| AL | Octopus | 10,769,198 | 1,980,872 | 12,750,070 | 84.5% |

## Discussion

We have shown that genotyping is substantially more error-prone in polyploids than in diploids using typical whole-genome sequencing depths, emphasising that caution must be used when interpreting polyploid genotype calls. We also found considerable differences in accuracy between callers. Notably, Octopus produced less than a quarter of the total errors of other methods, and half the errors on some datasets. We believe this is primarily due to Octopus modelling longer haplotypes during genotyping, as this increases statistical confidence in allele copy-number. Regenotyping with longer haplotypes increased genotyping precision, supporting this hypothesis.

Analysis of real autotriploid banana datasets revealed high discordance between callers, and more alarmingly, high discordance for callers on similar datasets of an identical specimen.

While these results were at least consistent with the relative accuracy of callers determined by our benchmarks using synthetic polyploid data (Octopus was the most concordant caller), absolute error rates were evidently higher in real polyploid data. Reasons for this may include: i) greater divergence from the reference genome^32^; ii) higher levels of repetitive elements in the genome^10^; iii) more structural variation^32^; iv) less complete reference genome^10^; v) higher rates of sequencing related errors, such as due to the use of PCR amplification. vi) bioinformatics algorithms optimised for human data.

Manual review of discordant banana calls using haplotype-tagged and realigned evidence BAMs generated by Octopus indicated that a large fraction were due to slightly different indels called in each dataset, suggesting failure to discover correct allele(s) in one or both datasets. Other indicative sources of error included lack of read depth, allele coverage bias, and reads with low mapping quality. These observations, which are consistent with the problems outlined above, could potentially be overcome with better read mapping^33^ and variant discovery methods, or long-read sequencing^34^.

We have only considered single sample polyploid calling in this work, however, multi-sample calling is important for studying population diversity. Population calling in humans is a difficult problem due to the computational complexities of joint calling and difficulties in merging independent callsets. Population calling in polyploids will likely be even more challenging, and would perhaps benefit from more sophisticated genotype prior models^23^.

Moving forward, there is clearly room for improvement in polyploid genotyping from sequencing. The creation of high quality validation sets with real polyploid samples would highly valuable in the development of polyploid calling algorithms, including Octopus. We hope that this work lays the groundwork for future developments.

## Acknowledgements

This work was supported by The Wellcome Trust Genomic Medicine and Statistics PhD Program (grant nos. 203735/Z/16/Z to D.P.C). The computational aspects of this research were supported by the Wellcome Trust Core Award Grant Number 203141/Z/16/Z and the NIHR Oxford BRC. The views expressed are those of the author(s) and not necessarily those of the NHS, the NIHR or the Department of Health. We would like to thank Len Trigg at Real Time Genomics for kindly arranging an update to RTG Tools for polyploid genotype support on our request.

## Author contributions

D.P.C formulated and did the analysis and wrote the paper. D.W. and G.L critically reviewed the manuscript and supervised the project.

## Ethics declaration

The authors declare no competing interests.

## Methods

### Synthetic polyploids with real reads

Raw reads (FASTQ) generated for the PrecisionFDA Truth v2 challenge^35^ were downloaded from the DNAnexus portal (https://precision.fda.gov/challenges/10). Each FASTQ was line counted to ensure realistic haplotype frequencies, before concatenating contributing samples to make the full data polyploid dataset. Downsampling was performed directly on the FASTQ files using *seqtk* with default seed. The sampling fraction was set using: *test depth* / *full depth*, where full depth is 35 × *ploidy*/2. Reads were mapped with BWA-MEM using default alignment parameters.

### Changes to Octopus for polyploid calling

While the models that we previously described for Octopus^17^ are fully capable of polyploid calls, in practice we found some issues. Runtimes were prohibiting for high ploidies due to the model always considering every possible genotype for a given set of candidate haplotypes, which is reasonable for diploids but not polyploids. Moreover, sensitivity for low copy-number variants was not optimal due to the variant discovery mechanisms not fully accounting for ploidy.

To resolve the runtime issue, we modified the genotype proposal algorithm so that an upper bound on the number of genotypes evaluated can be specified. The algorithm respects this limit by evaluating the full model on the maximum ploidy that results in less candidate genotypes than the limit for a given set of haplotypes, and then extends a subset of these with greatest posterior probability using each of the candidate haplotypes. The procedure is then applied iteratively, increasing the ploidy by one each iteration, until the desired ploidy is reached. We expect this procedure to work well when the number of unique haplotypes present in a region is not substantially greater than the first ploidy considered. We addressed the sensitivity issue by tweaking the pileup and local *de novo* reassembly candidate variant discovery algorithms to account for sample ploidy.

### Variant calling polyploids

For GATK4, we called variants using BAMs with marked duplicates created by GATK4’s *MarkDuplicates* tool. Raw BAMs were used for FreeBayes and Octopus. The sample ploidy was specified for all callers: --*organism-ploidy* (Octopus), --*sample-ploidy* (GATK4), and --*ploidy* (FreeBayes). For FreeBayes, we requested genotype qualities with the *-=* option.

### Filtering variant calls

For GATK4, we used filter expressions: “*filter ‘QD < 2.0’ –filter-name ‘QD2’ -filter ‘QUAL < 50’ –filter-name ‘Q50’ -filter ‘GQ < 5’ –filter-name ‘GQ5’ -filter ‘FS > 60.0’ –filter-name ‘FS60’ -filter ‘SOR > 3.0’ –filter-name ‘SOR3’ -filter ‘MQ < 40.0’ –filter-name ‘MQ40’ -filter ‘MQRankSum < −12.5’ –filter-name ‘MQRankSum-12.5’ -filter ‘ReadPosRankSum < −8.0’ –filter-name ‘ReadPosRankSum-8*’”. For FreeBayes, we used filter expression: “*QUAL > 1 & GQ > 1 & SAF > 0 & SAR > 0*”.

### Genotype and allele comparisons

We used RTG Tools vcfeval (v3.12) for genotype and allele comparisons, using the --*sample-ploidy* and --*ref-overlap* options. For allele matching, we also used the --*squash-ploidy, --XXcom.rtg.vcf.eval.flag-alternates=true*, and --*output-mode=“annotate*” options, and then determined true and false calls based on the resulting BASE, CALL, BASE_ALTERNATE, and CALL_ALTERNATE annotations.

### Identifying copy-number errors

Biallelic copy-number errors were identified by running RTG Tools vcfeval with the --*output-mode=“combine*” option, and considering biallelic calls with baseline INFO annotations “BASE=FN_CA” and “CALL=FP_CA”.

### Long haplotypes with Octopus

To call long haplotypes with Octopus, we provided Octopus with the variant calls it previously produced with default setting as canidates (--*sourcecandidates*) and disabled *de novo* variant discovery (--*disable-denovo-variant-discovery*). We also set command line options --*lagging-level=OPTIMSITIC, --backtrack-level=AGGRESSIVE*, and --*max-haplotypes=400*.

### Banana concordance analysis

Callsets for the banana datasets were intersected using a custom script (https://github.com/dancooke/starfish) that invokes both RTG Tools vcfeval (that only supports 2-way comparisons) and bcftools to achieve multisample haplotype-aware comparisons. UpSet plots were created with UpSetR^36^.

## Code availability

Octopus source code and documentation is freely available under the MIT licence from https://github.com/luntergroup/octopus. Custom Snakemake and Python code used for data analysis is available from https://github.com/luntergroup/polyploid.

## Data availability

All raw sequencing data used for the synthetic polyploid benchmarks was previously published in the Precision FDA Truth Challenge v235; synthetic datasets can easily be reproduced from the raw data using the online code. The *Musa acuminata* sequencing data has been published previously^32^.

## Notes

### Competing Interest Statement

The authors have declared no competing interest.

