## Supplementary Material for "Benchmarking small-variant genotyping in polyploids"

### Benchmarking small-variant genotyping in polyploids: supplementary material

#### 1 Supplementary Tables and Figures

**(a)**

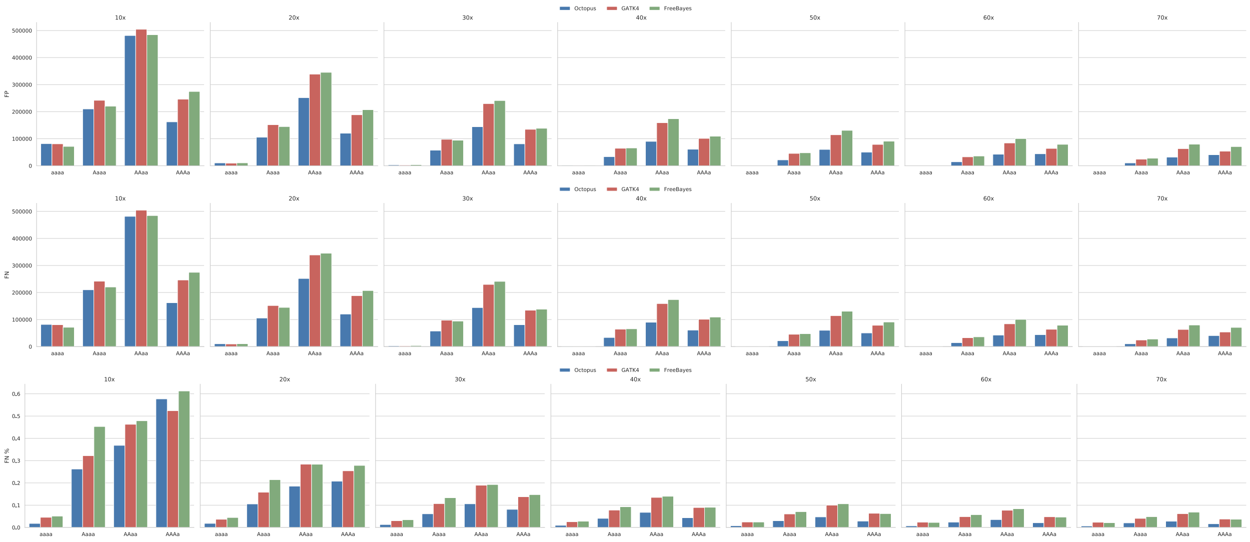


**(b)**

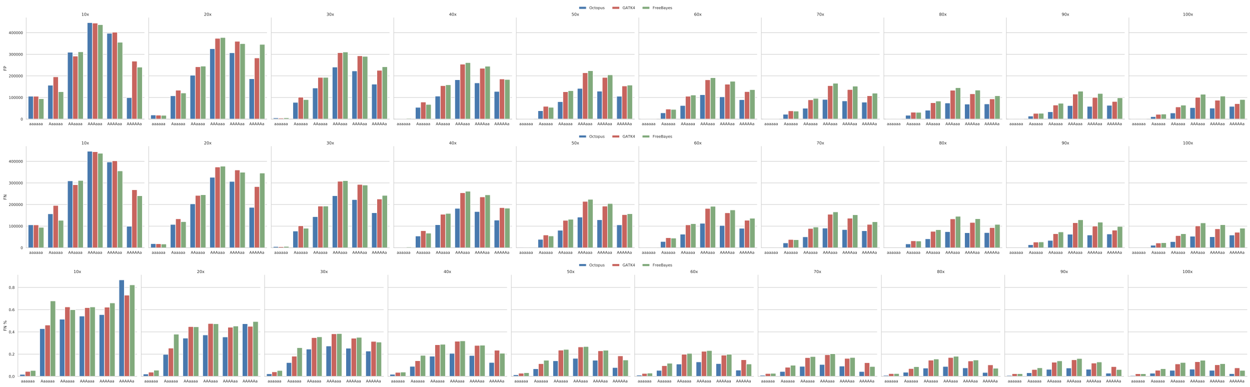


**Supplementary Figure. 1 — Biallelic genotyping errors in synthetic polyploid samples. a** Tetraploid. **b** Hexaploid.

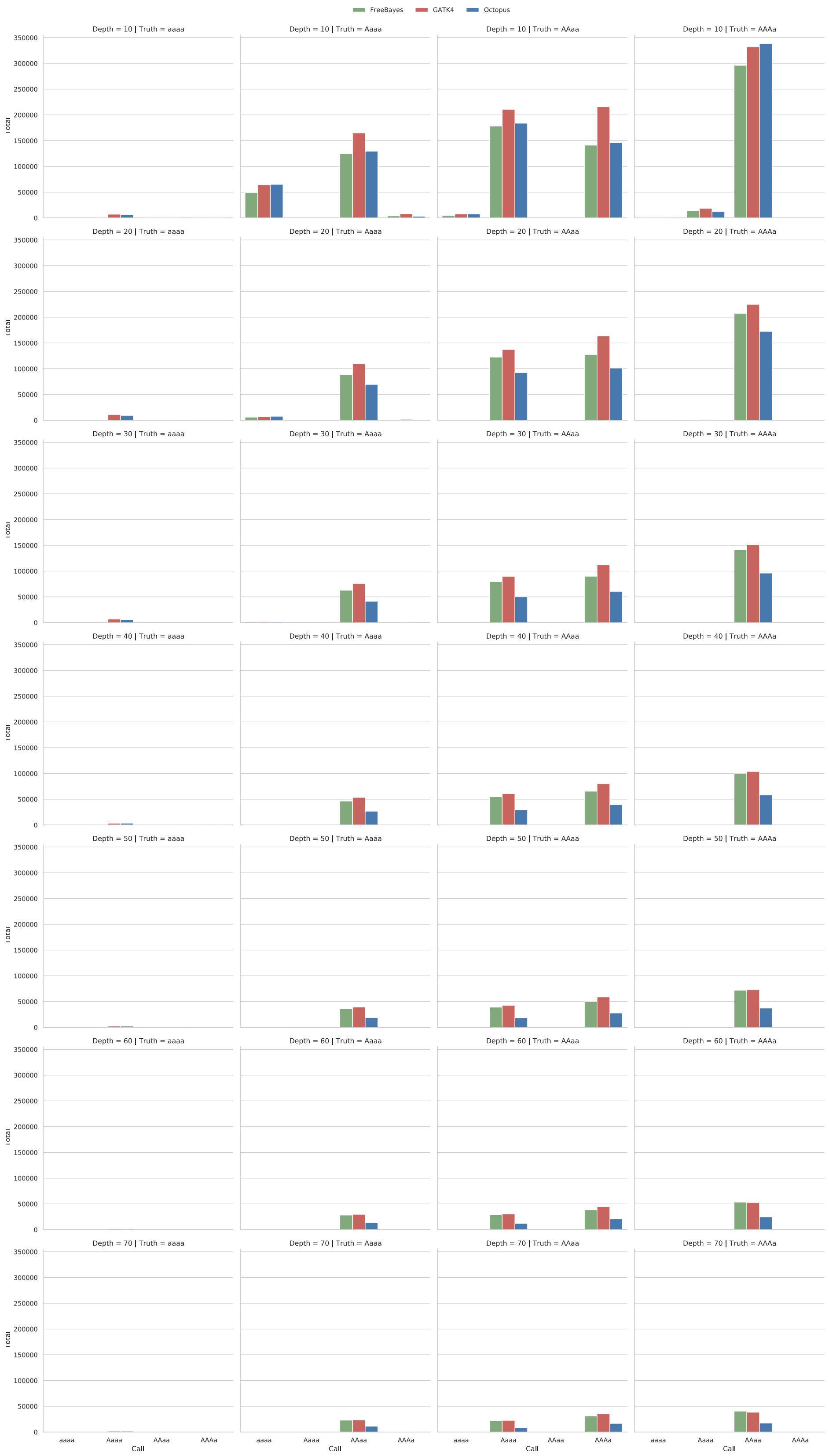


**Supplementary Figure. 2 — Biallelic copy number errors in synthetic tetraploid samples.**

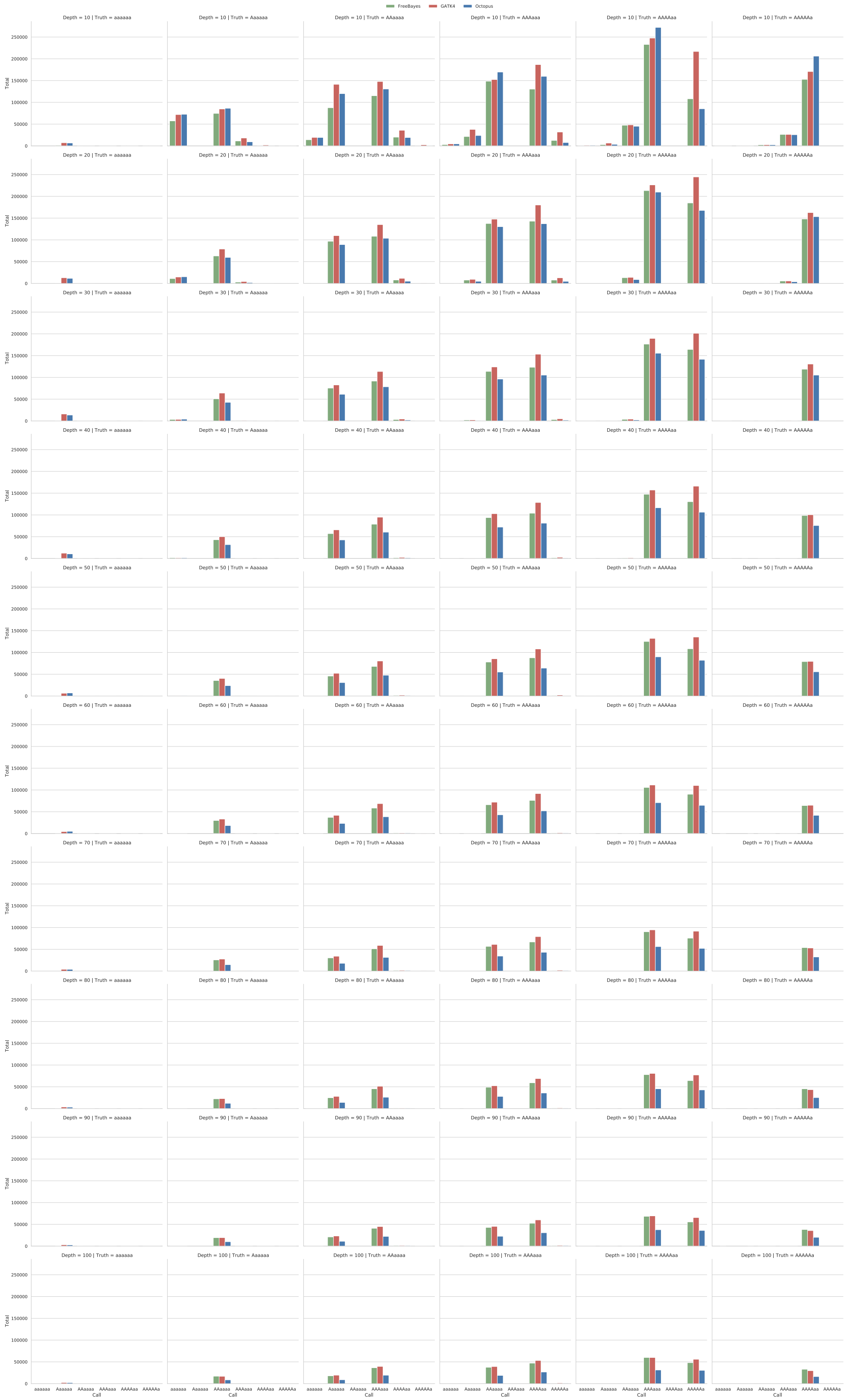


**Supplementary Figure. 3 — Biallelic copy number errors in synthetic hexaploid samples.**

(a)

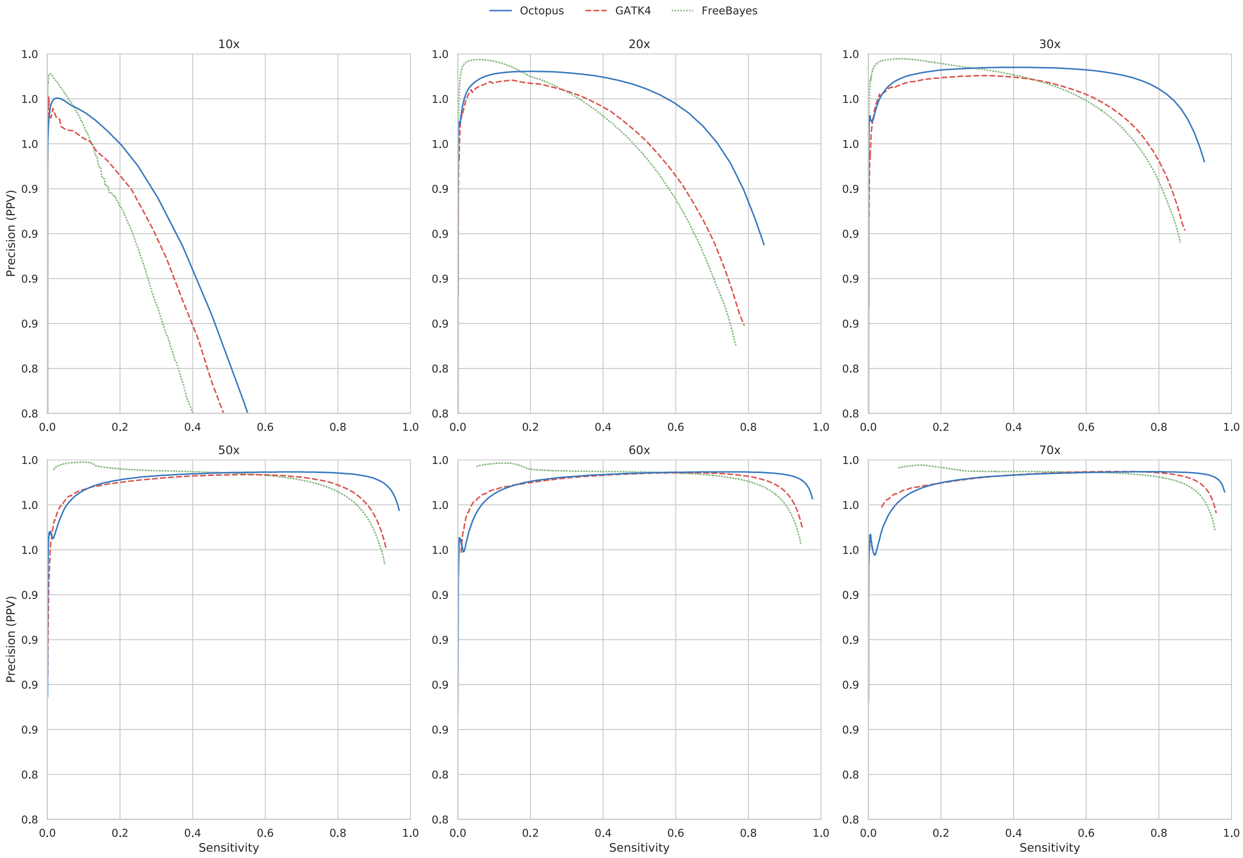


(b)

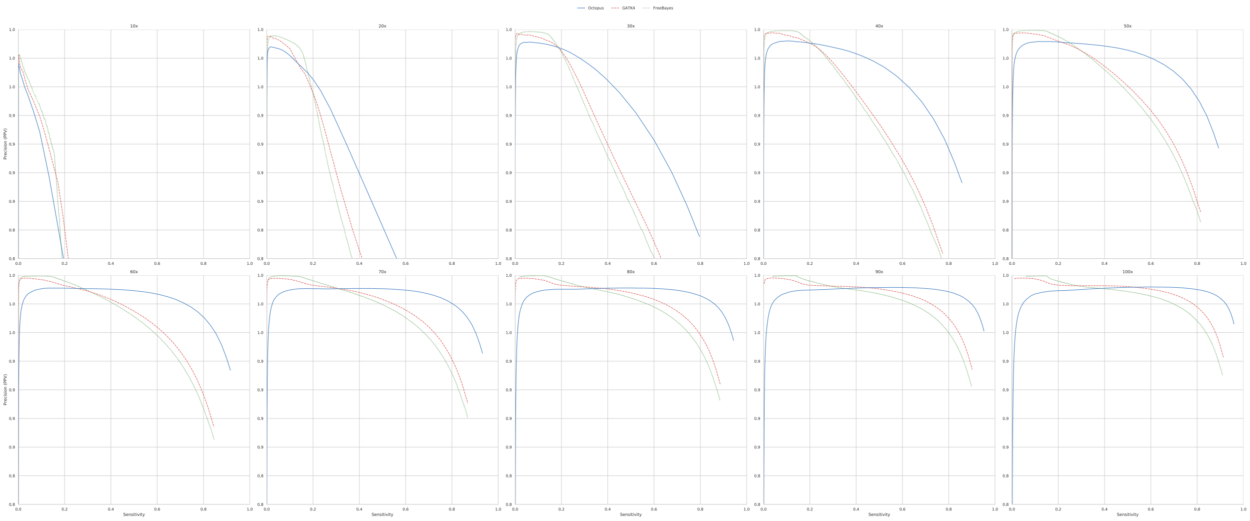

**Supplementary Figure. 4 — Precision-recall curves for genotyping errors in synthetic polyploid samples.** GQ was used to generate curves for all samples. **a** Tetraploid. **b** Hexaploid.

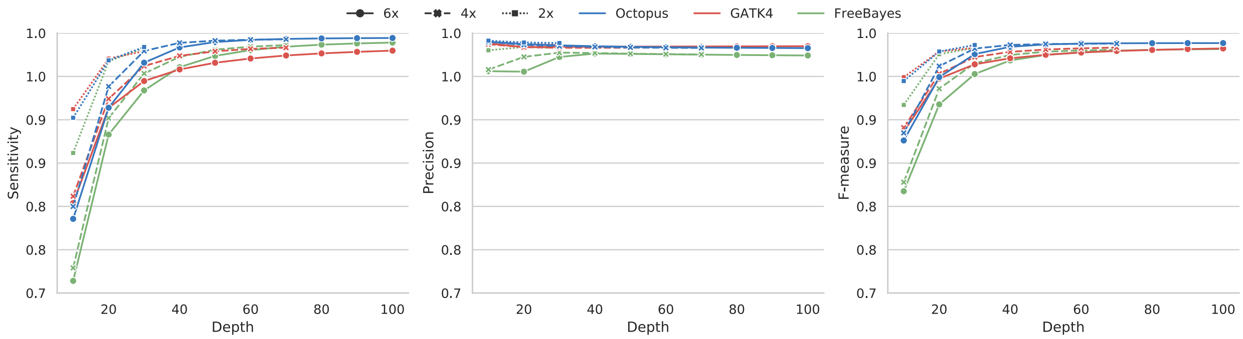


**Supplementary Figure. 5 — Comparison of allele calling accuracies for diploid and synthetic polyploid datasets**.

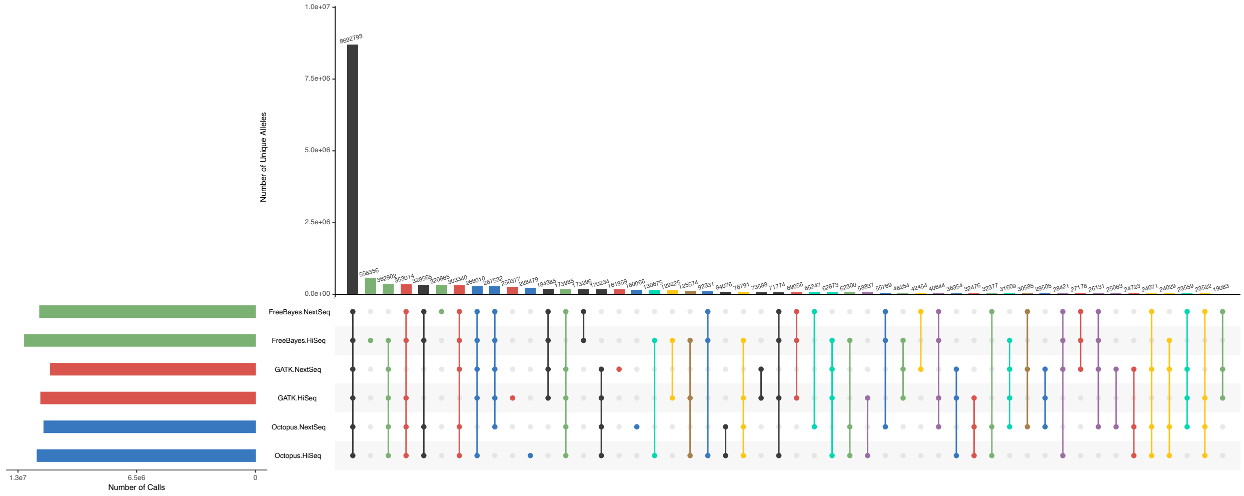


**Supplementary Figure. 6 — Comparison of alleles called in two Illumina datasets (HiSeq and NextSeq) of banana specimen by Octopus, GATK4, and FreeBayes**. UpSet plot shows callset intersections for each caller-dataset pair. The largest 50/63 intersection sets are shown. Intersections are color coded by caller discordance between the two datasets: No discordances (black), Octopus (blue), GATK4 (red), FreeBayes (green), Octopus & GATK4 (purple), Octopus & FreeBayes (cyan), GATK4 & FreeBayes (yellow), All (brown). The total number of unique alleles calls was 14,825,250.
